# Association of whole-genome and NETRIN1 signaling pathway-derived polygenic risk scores for Major Depressive Disorder and thalamic radiation white matter microstructure in UK Biobank

**DOI:** 10.1101/282053

**Authors:** Miruna C. Barbu, Yanni Zeng, Xueyi Shen, Simon R. Cox, Toni-Kim Clarke, Jude Gibson, Mark J. Adams, Mandy Johnstone, Chris S. Haley, Stephen M. Lawrie, Ian J. Deary, Major Depressive Disorder Working Group of the Psychiatric Genomics Consortium, 23andMe Research Team, Andrew M. McIntosh, Heather C. Whalley

## Abstract

**Background:** Major Depressive Disorder (MDD) is a clinically heterogeneous psychiatric disorder with a polygenic architecture. Genome-wide association studies have identified a number of risk-associated variants across the genome, and growing evidence of NETRIN1 pathway involvement. Stratifying disease risk by genetic variation within the NETRIN1 pathway may provide an important route for identification of disease mechanisms by focusing on a specific process excluding heterogeneous risk-associated variation in other pathways. Here, we sought to investigate whether MDD polygenic risk scores derived from the NETRIN1 signaling pathway (NETRIN1-PRS) and the whole genome excluding NETRIN1 pathway genes (genomic-PRS) were associated with white matter integrity.

**Methods:** We used two diffusion tensor imaging measures, fractional anisotropy (FA) and mean diffusivity (MD), in the most up-to-date UK Biobank neuroimaging data release (FA: N = 6,401; MD: N = 6,390).

**Results:** We found significantly lower FA in the superior longitudinal fasciculus (β = -0.035, p_corrected_ = 0.029) and significantly higher MD in a global measure of thalamic radiations (β = 0.029, p_corrected_ = 0.021), as well as higher MD in the superior (β = 0.034, p_corrected_ = 0.039) and inferior (β = 0.029, p_corrected_ = 0.043) longitudinal fasciculus and in the anterior (β = 0.025, p_corrected_ = 0.046) and superior (β = 0.027, p_corrected_ = 0.043) thalamic radiation associated with NETRIN1-PRS. Genomic-PRS was also associated with lower FA and higher MD in several tracts.

**Conclusions:** Our findings indicate that variation in the NETRIN1 signaling pathway may confer risk for MDD through effects on thalamic radiation white matter microstructure.

## Introduction

Major Depressive Disorder (MDD) is a common and frequently disabling psychiatric disorder and a leading cause of disability worldwide (1). MDD is known to result from a complex combination of environmental and genetic factors (2; 3), with a moderate heritability of approximately 37% (4; 5; 6).

Genome-wide association studies (GWAS) suggest that at least part of MDD‘s heritability is due to the cumulative effect of alleles of small effect size (7; 8) and have identified a number of risk-associated genetic variants across the genome (6; 7; 9; 10; 11). Significant findings for GWAS analyses can also be annotated to specific biological pathways, revealing underlying cellular and molecular mechanisms.

Following several GWAS, the Psychiatric Genomics Consortium (PGC) have identified an aggregation of variants in several specific biological pathways (12; 13). In MDD, Zeng et al. (2017) (14) combined pathway analysis and regional heritability analysis in two independent samples and reported that the NETRIN1 signaling pathway was involved in the genetic aetiology of MDD. Moreover, polygenic risk scores (PRS) calculated for this pathway alone more accurately predicted MDD in one of the cohorts compared to PRS calculated for the whole genome. Genetic variation within the NETRIN1 signaling pathway may therefore capture more aetiologically circumscribed liability for MDD that is less susceptible to heterogeneous influences from other biological pathways.

Animal studies have previously indicated that NETRIN1, by binding to and activating NETRIN1 receptors such as ‘Deleted in Colorectal Cancer’ (DCC), plays an important role in commissural and cortical axon guidance (15). More recently, DCC was identified as playing a crucial role in thalamic axonal growth, confirming that interaction of NETRIN1 with DCC leads to successful axon growth during central nervous system development (16). GWAS of other traits related to MDD have also shown an aggregation of variants in the NETRIN1 pathway (17; 18).

Previous studies have attempted to investigate psychiatric disorders by examining relevant quantitative traits such as brain structure or function (19). Differences in white matter (WM) integrity as measured by diffusion tensor imaging (DTI) have been found between MDD patients and healthy participants in numerous studies, although findings have been widely inconsistent (20; 21; 22). For example, Shen et al. (2017) (20) found significantly lower global white matter integrity in association fibres and thalamic radiations, as measured by fractional anisotropy (FA), in MDD patients compared to healthy individuals. More specifically, they also found lower FA in the left superior longitudinal fasciculus, superior thalamic radiations and forceps major tracts in MDD patients. Lower WM integrity as measured by FA has also been found in adolescents with MDD as compared to age-matched healthy individuals (21; 22).

It has previously been shown that the NETRIN1 signaling pathway is associated with MDD and white matter microstructure (14). Therefore, in the current study, we sought to investigate the association between MDD risk-associated variants in the NETRIN1 signaling pathway and white matter integrity. We first created polygenic risk scores for pathway SNPs (NETRIN1-PRS) and SNPs excluded from the pathway (genomic-PRS). We then tested their association with WM integrity as measured by FA and mean diffusivity (MD). We used the most up-to-date genetic (N = 488,366) and imaging (N DTI = 8,839) data available from UK Biobank (UKB). We hypothesized that NETRIN1-PRS would be significantly associated with WM integrity, after adjustment for genomic-PRS, indicating a potential role of the pathway in MDD pathophysiology.

## Methods and Materials

### UK Biobank

The UKB study consists of 502,617 community-dwelling individuals who were recruited between 2006 and 2010 in the United Kingdom (http://biobank.ctsu.ox.ac.uk/crystal/field.cgi?id=200). UKB received ethical approval from the Research Ethics Committee (reference: 11/NW/0382). This study has been approved by the UKB Access Committee (Project #4844). Written informed consent was obtained from all participants.

### Study population

In the most recent UKB imaging data release, 8,839 individuals (N female = 4,639; N male = 4,200; mean age: 62.54 +/- 7.42 years; age range: 45.17 – 79.33) completed DTI assessment, and a quality check by UKB. In addition to this, for the current study, individuals were excluded if they participated in studies from the PGC MDD GWAS (24) or Generation Scotland (Scottish Family Health Study), or if they happened to be related, as the PGC MDD GWAS dataset was used in order to calculate PRS. Moreover, individuals whose FA and MD values were greater than three standard deviations above/below the mean were not included in the study (Supplementary Material, tables S4 and S5). This resulted in 6,401 individuals with FA values (N female = 3,334; N male = 3,067; mean age: 62.60 +/- 7.37; age range: 45.92 – 78.42) and 6,390 individuals with MD values (N female = 3,327; N male = 3,063; mean age: 62.58 +/- 7.36; age range: 45.92 – 78.42), excluding 19 and 30 individuals with FA and MD values from a total of 6,420, respectively. Details of data exclusion as well as participant information for the full dataset (N = 6,420) are shown in the Supplementary Material (tables S1 and S2).

### SNP annotation

Genic SNPs found in the NETRIN1 signaling pathway as taken from Zeng et al.‘s (2017) study (14) (N genes = 43; gene list is presented in the Supplementary Material, table S3) and genic SNPs excluded from the pathway were annotated using the program ANNOVAR. ANNOVAR is a biostatistical tool used to annotate genetic variants to functional genomic regions (genome build used in current study: hg 19) (23). In the current study, we performed a gene-based annotation for SNPs used in the largest available GWAS of MDD (N=461,134, of which 130,664 were MDD cases), carried out by the Psychiatric Genomics Consortium (24), which includes summary statistics from the personal genetics company 23andMe, Inc. (10). We defined gene boundaries as an extended region of 20 kb from transcription start sites and transcription end sites. After SNPs were annotated to genes, they were further mapped to the NETRIN1 signalling pathway. All protein-coding genes within this file were annotated in reference to hg 19. Intergenic SNPs were not included in the annotated files. The resulting output file included: function of each SNP, gene name, chromosome number, start position, end position, reference and alternative alleles, odds ratio, standard error and p-value for each variant.

Following functional annotation, a file containing the 43 gene names included in the NETRIN1 signaling pathway was used as an input in order to extract gene-based SNPs located in the pathway. For the genomic-PRS, all gene-based SNPs excluding those implicated in the NETRIN1 signaling pathway were extracted. The two files were then used as input for creation of PRS.

### Genotyping and PRS profiling

A total of 488,363 UKB blood samples (N female = 264,857; N male = 223,506; http://biobank.ctsu.ox.ac.uk/crystal/field.cgi?id=22001), were genotyped using two different arrays: UK BiLEVE array (N = 49,949) (http://biobank.ctsu.ox.ac.uk/crystal/refer.cgi?id=149600) and UK Biobank Axiom array (N = 438,417) (http://biobank.ctsu.ox.ac.uk/crystal/refer.cgi?id=149601). Details of genotyping and quality control are described in more detail by Hagenaars et al. (2016) (25) and Bycroft et al. (2017) (26).

Using the largest available GWAS of MDD, PRS for each individual were computed using PRSice (27), at five p-value thresholds (0.01, 0.05, 0.1, 0.5, 1) by adding the number of risk alleles and weighting them by the strength of association with MDD. PRS were created both from SNPs annotated to the NETRIN1 signalling pathway and from SNPs from the rest of the genome, thus resulting in separate PRS lists. PRS were created both with and without clump-based pruning of SNPs in linkage disequilibrium (r2 = 0.25, 250km window). The primary analysis reported in this manuscript concerns unpruned SNPs, owing to the potential of causal variants within the NETRIN1 pathway to be in LD with other variants, and uses SNPs which met a significance level of p = 0.5, in line with previous studies (28; 29). Secondary analyses with other PRS p-value thresholds, as well as with LD pruned SNPs, are presented in the Supplementary Material (Tables S6 – S21).

### MRI acquisition

In the present study, imaging-derived phenotypes (IDPs) produced by UKB were used. MRI acquisition and pre-processing procedures for FA and MD values of white matter tracts were performed by UKB using standardised protocols (https://biobank.ctsu.ox.ac.uk/crystal/docs/brain_mri.pdf). Briefly, images were collected on a single Siemens Skyra 3.0 T scanner with a standard Siemens 32-channel head coil and were pre-processed using FSL packages; parcellation of white matter tracts was conducted using AutoPtx (30).

Summary data were composed of tract-averaged FA and MD values for 15 major white matter tracts, of which 12 are bilateral and three are unilateral. The white matter tracts were also categorised into three separate subsets, as follows: association fibres: inferior fronto-occipital fasciculus, uncinate fasciculus, cingulum bundle (gyrus and parahippocampal), superior and inferior longitudinal fasciculus; thalamic radiation fibres: anterior, superior and posterior thalamic radiations; projection fibres: forceps major and minor, corticospinal tract, acoustic radiation, medial lemniscus and middle cerebellar peduncle. Global measures of FA and MD are referred to as general factors of FA and MD (gFA and gMD, respectively).

Exclusion criteria comprised removal of scans with severe normalisation problems by UKB. Moreover, individuals whose FA and MD values were higher than three standard deviations from the sample mean were also excluded. Results for the full dataset with outliers included are also presented in the Supplementary Material (tables S1 and S2). Lastly, due to the fact that the position of the head and radio-frequency coil in the scanner may affect data quality as well as IDPs, three scanner brain position variables which may be used as confounding variables in subsequent analyses were generated by UKB: lateral brain position – X (http://biobank.ctsu.ox.ac.uk/crystal/field.cgi?id=25756), transverse brain position –Y (http://biobank.ctsu.ox.ac.uk/crystal/field.cgi?id=25757) and longitudinal brain position – Z (http://biobank.ctsu.ox.ac.uk/crystal/field.cgi?id=25758). The three variables were included as covariates in the statistical analysis described below.

### Statistical methods

All analyses were conducted using R (version 3.2.3) in a Linux environment. In order to test the association between the NETRIN1 signaling pathway- and genomic pathway-derived unpruned PRS lists, we used repeated measures linear mixed-effects models (function “lme” in package “nlme”) for 12 bilateral brain regions, correcting for hemisphere, with age, age^2^, sex, fifteen genetic principal components, three MRI head position coordinates and genotype array set as covariates. For unilateral tracts, global measures of FA and MD, and tract categories, we used a general linear model (function “lm”), using the same covariates as above, and without hemisphere included as a separate term in the model. All models included both the genomic-PRS and the NETRIN1-PRS as predictor variables.

First, we tested the association between unpruned PRS (both NETRIN1-PRS and genomic-PRS) and global white matter integrity. We applied principal component analysis (PCA) on the 27 white matter tracts (12 tracts in both the right and left hemisphere and three unilateral tracts) in order to extract a latent measure. Scores of the first unrotated component of FA and MD (variance explained = 37.52% for FA and 38.83% for MD) were extracted and set as the dependent variable in a general linear model in order to test association with both NETRIN1-PRS and genomic-PRS.

We then examined the three categories of white matter tracts by applying PCA on the regions involved in each, as a substantial proportion of white matter microstructural properties shows substantial commonality across these pathways (31). Scores of the first unrotated component of FA and MD were similarly extracted and set as dependent variables in general linear modelling, as above. Variance explained for each white matter tract subset was as follows: association fibres: 45.36% (FA), 50.76% (MD); thalamic radiations: 60.85% (FA), 73.40% (MD); projection fibres: 35.54% (FA), 29.28% (MD).

Lastly, we tested the association between PRS (both NETRIN1-PRS and genomic-PRS) and each individual white matter tract (N = 15). We used a repeated-effect linear model for the 12 bilateral tracts and a random-effect general linear model for the three unilateral tracts. False discovery rate correction was applied over all tracts.

### Permutation analysis

In order to establish that the effect of the NETRIN1 pathway-derived PRS on WM integrity as measured by FA and MD was real and not due to chance, a circular genomic permutation was applied to the pathway SNP genotypes (32). This was done by creating PRS from 1000 circularly permuted SNP lists with the same set size as the NETRIN1 pathway. The 1000 PRS lists were then fitted in linear mixed-effects and general linear models, depending on the white matter tract tested, and their association with five white matter tracts and one tract category, found to be significantly associated with NETRIN1, was tested.

## Results

Results presented below are significant specifically to each pathway. White matter tracts showing a significant association with both the NETRIN1-PRS and the genomic-PRS pathways are described in the supplementary materials. Results for all individual white matter tracts, tract categories and global measures can be found in tables 1-4 and figures 1-4.

**Table 1.**
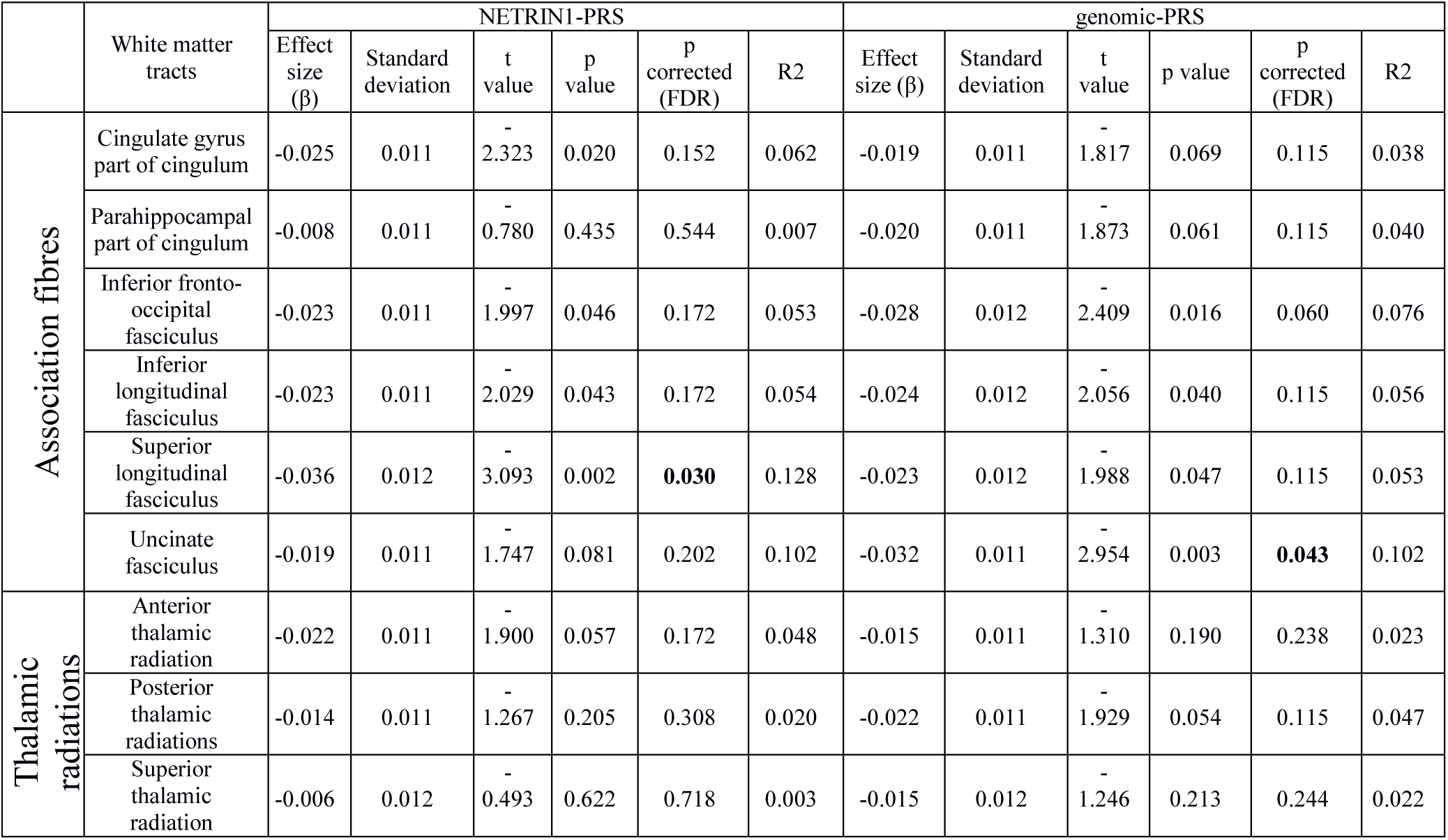
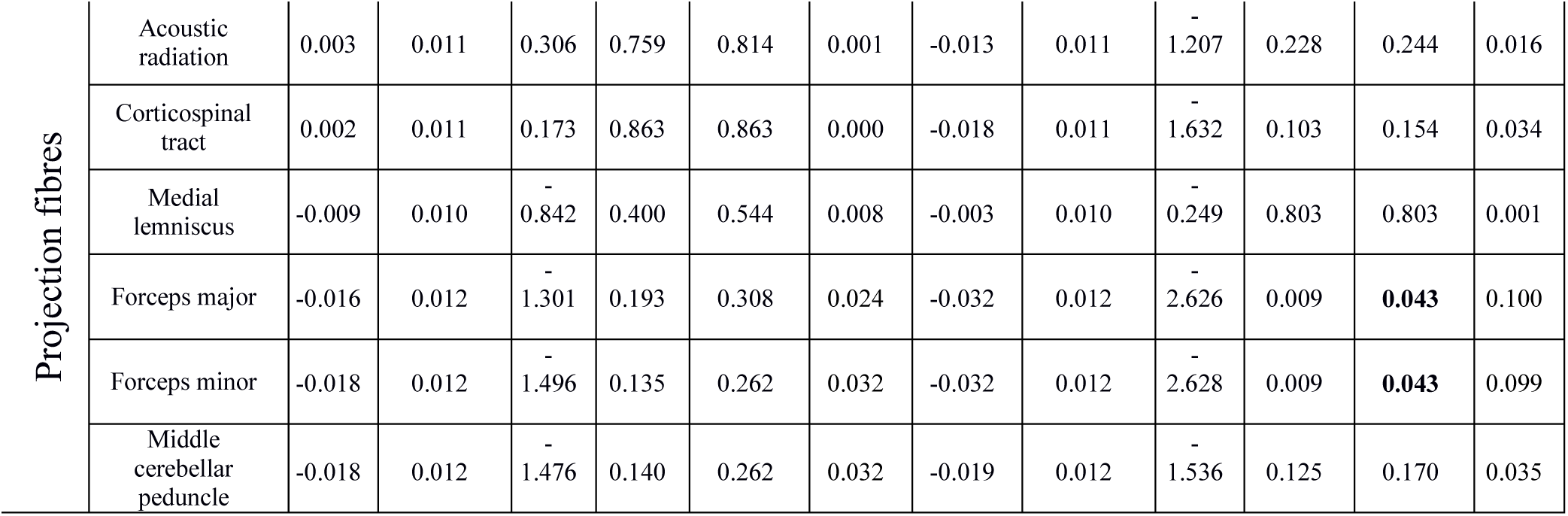
The effect of NETRIN1-PRS & genomic-PRS at PRS threshold 0.5 on individual white matter tracts (FA values). Statistically significant p-values after false discovery rate correction for each pathway individually are shown in bold. R2 = estimate of variance explained by each pathway in %.

**Table 2.** The effect of NETRIN1-PRS & genomic-PRS at PRS threshold 0.5 on global FA and 3 white matter tract categories. Statistically significant p-values after false discovery rate correction for each pathway individually are shown in bold. R2 = estimate of variance explained by each pathway in %.

|  | NETRIN1-PRS |  |  |  |  |  | genomic-PRS |  |  |  |  |  |
| --- | --- | --- | --- | --- | --- | --- | --- | --- | --- | --- | --- | --- |
| | Effect size($\beta$) | Standard deviation | t value | p value | p corrected (FDR) | R2 | Effect size($\beta$) | Standard deviation | t value | p value | p corrected (FDR) | R2 |
| gFA | -0.026 | 0.012 | -2.197 | 0.028 | 0.056 | 0.068 | -0.033 | 0.012 | -2.769 | 0.006 | **0.011** | 0.109 |
| Association fibres | -0.033 | 0.012 | -2.762 | 0.006 | 0.023 | 0.107 | -0.034 | 0.012 | -2.836 | 0.005 | 0.011 | 0.113 |
| Thalamic radiations | -0.018 | 0.012 | -1.482 | 0.138 | 0.185 | 0.032 | -0.022 | 0.012 | -1.855 | 0.064 | 0.064 | 0.050 |
| Projection fibres | -0.011 | 0.012 | -0.904 | 0.366 | 0.366 | 0.012 | -0.029 | 0.012 | -2.415 | 0.016 | **0.021** | 0.083 |

**Table 3.**
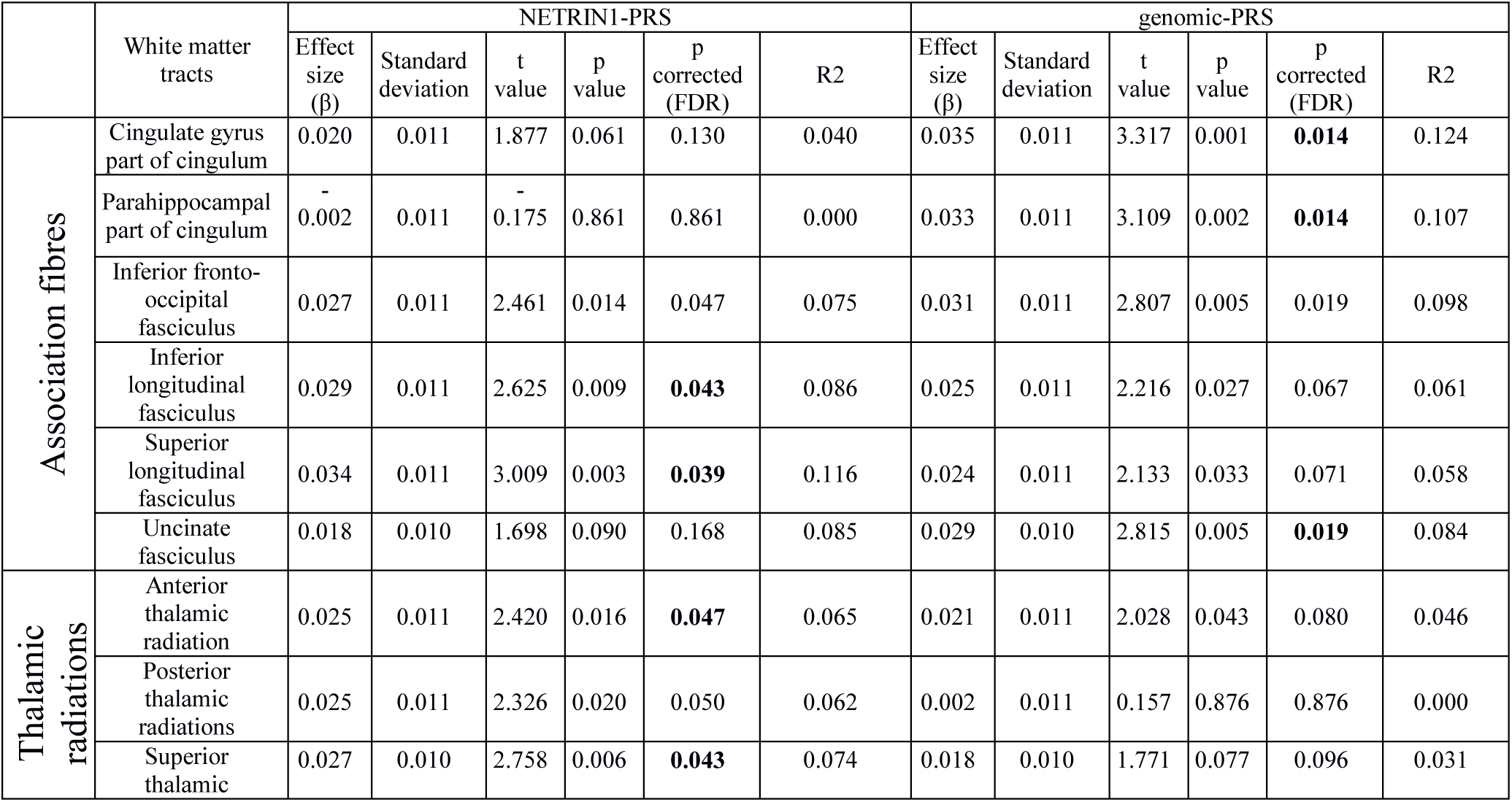
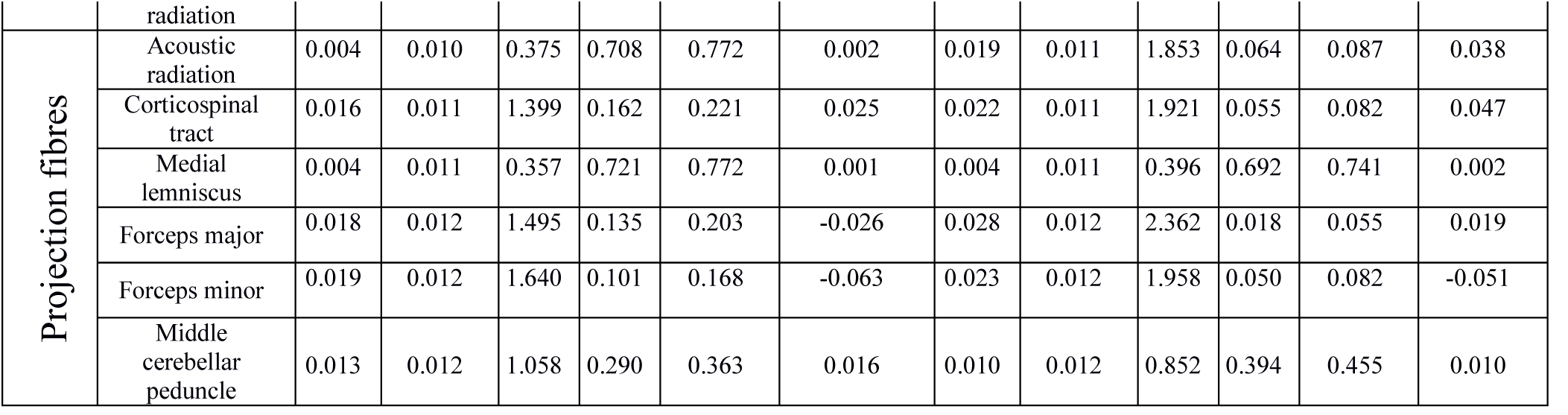
The effect of NETRIN1-PRS & genomic-PRS at PRS threshold 0.5 on individual white matter tracts (MD values). Statistically significant p-values after false discovery rate correction for each pathway individually are shown in bold. R2 = estimate of variance explained by each pathway in %.

**Table 4.** The effect of NETRIN1-PRS & genomic-PRS at PRS threshold 0.5 on global MD and 3 white matter tract subsets. Statistically significant p-values after false discovery rate correction for each pathway individually are shown in bold. R2 = estimate of variance explained by each pathway in %.

|  | NETRIN1-PRS |  |  |  |  |  | genomic-PRS |  |  |  |  |  |
| --- | --- | --- | --- | --- | --- | --- | --- | --- | --- | --- | --- | --- |
| | Effect size($\beta$) | Standard deviation | t value | p value | P corrected (FDR) | R2 | Effect size($\beta$) | Standard deviation | t value | p value | P corrected (FDR) | R2 |
| gMD | 0.028 | 0.011 | 2.417 | 0.016 | 0.031 | 0.076 | 0.034 | 0.011 | 2.924 | 0.003 | 0.007 | 0.111 |
| Association fibres | 0.022 | 0.012 | 1.897 | 0.058 | 0.077 | 0.048 | 0.042 | 0.012 | 3.591 | 0.000 | **0.001** | 0.172 |
| Thalamic radiations | 0.030 | 0.011 | 2.785 | 0.005 | **0.021** | 0.089 | 0.013 | 0.011 | 1.232 | 0.218 | 0.218 | 0.017 |
| Projection fibres | 0.021 | 0.012 | 1.766 | 0.077 | 0.077 | 0.045 | 0.029 | 0.012 | 2.380 | 0.017 | **0.023** | 0.081 |

**Figure 1.**
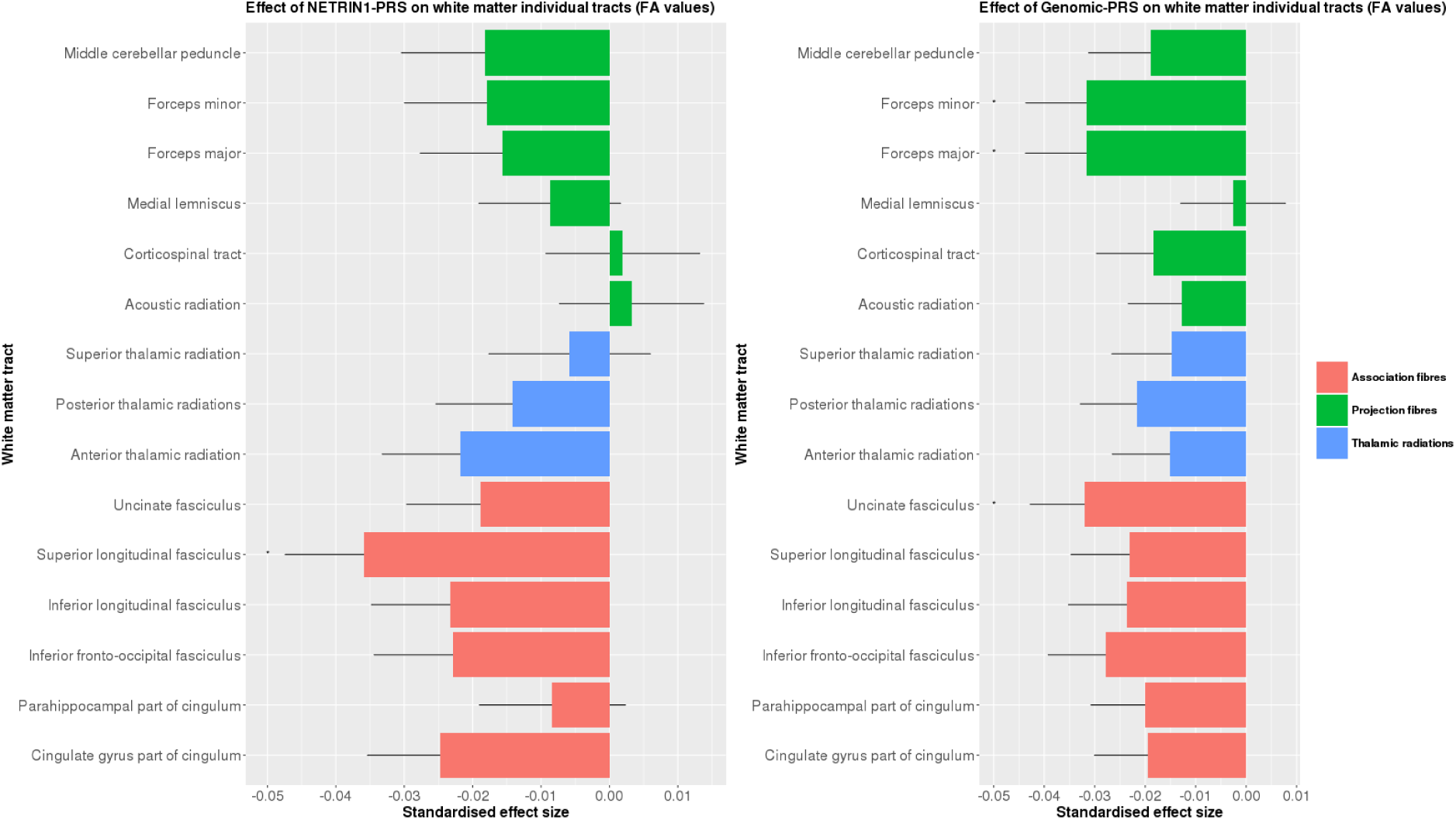
The effect of NETRIN1-PRS & genomic-PRS on FA values of white matter tracts. The x-axis indicates the standardised effect size of each pathway’s PRS and the y-axis indicates the white matter tracts. The legend indicates the tract category belonging to each white matter tract. The error bar represents standard deviation of mean.

**Figure 2.**
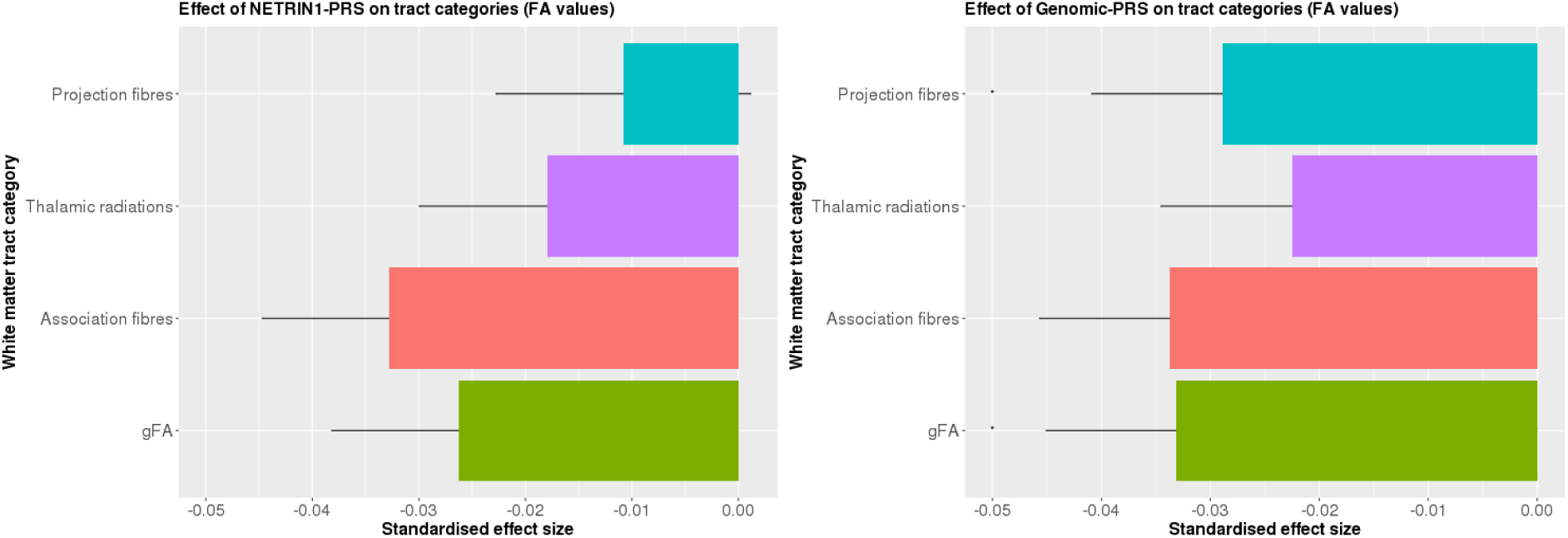
The effect of NETRIN1-PRS & genomic-PRS on FA values of tract categories and global FA. The x-axis indicates the standardised effect size of each pathway’s PRS and the y-axis indicates the tract categories. The error bar represents standard deviation of mean.

**Figure 3.**
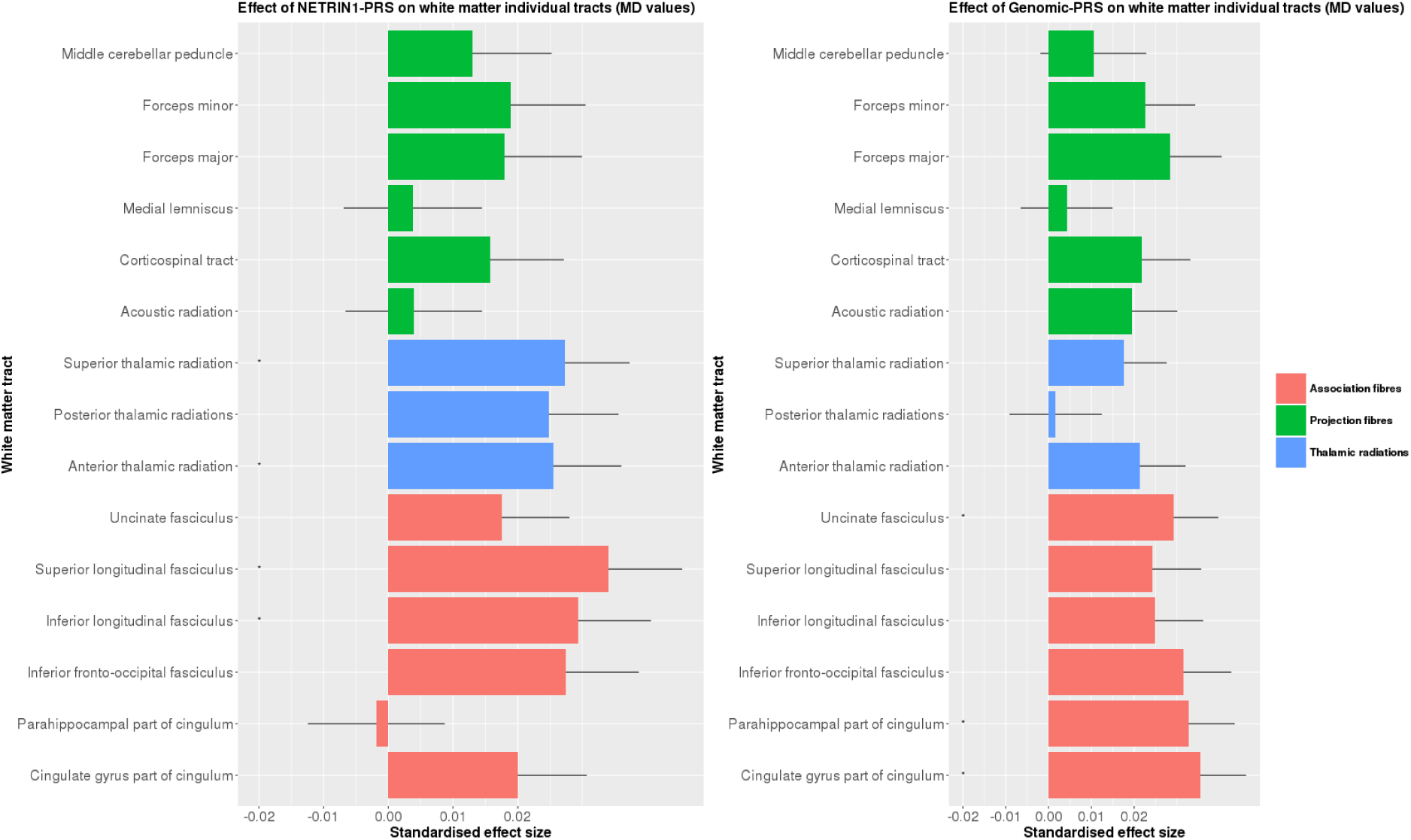
The effect of NETRIN1-PRS & genomic-PRS on MD values of white matter tracts. The x-axis indicates the standardised effect size of each pathway’s PRS and the y-axis indicates the white matter tracts. The legend indicates the tract category belonging to each white matter tract. The error bar represents standard deviation of mean.

**Figure 4.**
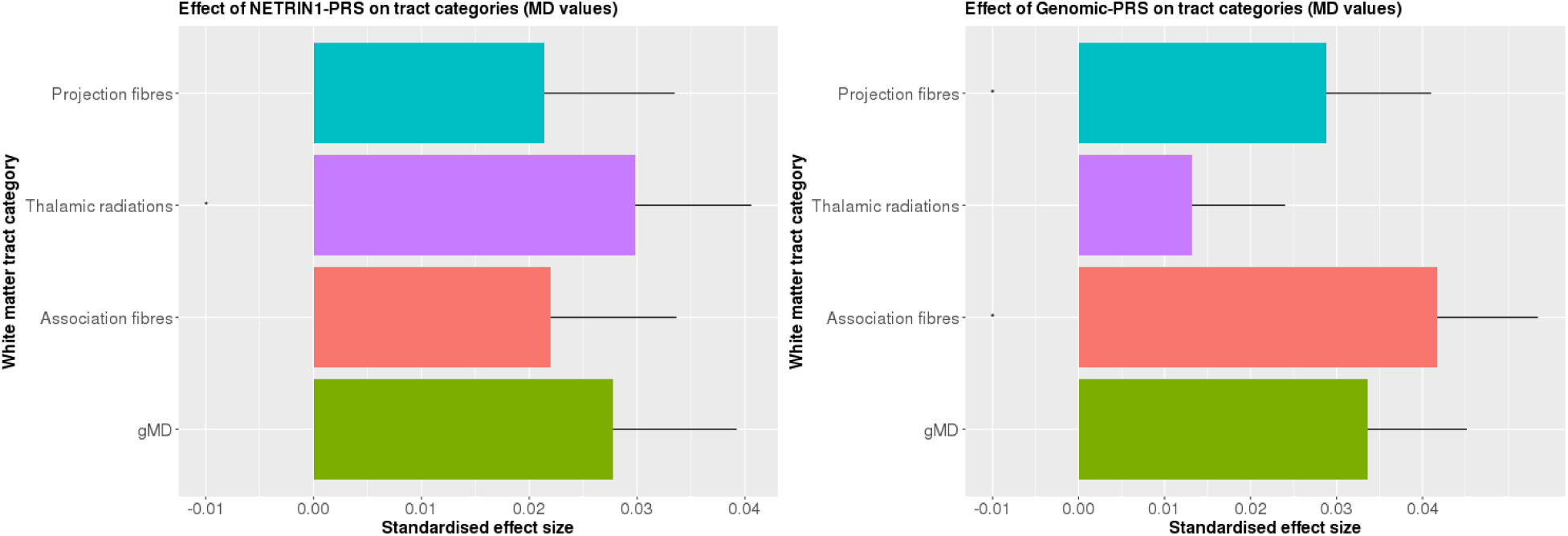
The effect of NETRIN1-PRS & genomic-PRS on MD values of tract categories and global MD. The x-axis indicates the standardised effect size of each pathway’s PRS and the y-axis indicates the tract categories. The error bar represents standard deviation of mean.

### The effect of unpruned NETRIN1-PRS & genomic-PRS on measures of white matter integrity – FA (N = 6,401)

#### Global measures

We first tested the effect of NETRIN1-PRS and genomic-PRS on global FA (gFA). Lower gFA was significantly associated with higher genomic-PRS (β = -0.033, p_corrected_ = 0.011).

#### Tract categories

We then tested the association between NETRIN1-PRS and Genomic-PRS and three subsets of white matter tracts (association fibres, thalamic radiations and projection fibres). Significantly lower values in projection fibres were found for genomic-PRS (β = -0.028, p_corrected_ = 0.020).

#### Individual white matter tracts

Lastly, we investigated the effect of NETRIN1-PRS and genomic-PRS on WM integrity in 15 individual white matter tracts. With regards to NETRIN1-PRS, we found significantly lower FA in the superior longitudinal fasciculus (β = -0.035, p_corrected_ = 0.029).

In the genomic-PRS, we found a significantly lower FA in the forceps major (β = -0.031, p_corrected_ = 0.043), forceps minor (β = -0.031, p_corrected_ = 0.043) and uncinate fasciculus (β = - 0.031, p_corrected_ = 0.043).

### The effect of unpruned NETRIN1-PRS & genomic-PRS on measures of white matter integrity – MD (N = 6,390)

#### Tract categories

MD values for association fibres (β = 0.041, p_corrected_ = 0.001) and projection fibres (β = 0.028, p_corrected_ = 0.023) were found to be significantly higher for genomic-PRS. MD values for thalamic radiations were found to be significantly higher in the NETRIN1-PRS (β = 0.029, p_corrected_ = 0.021).

#### Individual white matter tracts

Within the 15 individual white matter tracts, we found numerous areas significantly associated with both the NETRIN1-PRS and genomic-PRS. With regards to NETRIN1-PRS, MD values were significantly higher in the inferior longitudinal fasciculus (β = 0.029, p_corrected_ = 0.043), superior longitudinal fasciculus (β = 0.034, p_corrected_ = 0.039), and in the anterior (β = 0.025, p_corrected_ = 0.046) and superior (β = 0.027, p_corrected_ = 0.043) thalamic radiations.

In the genomic-PRS, we found significantly higher MD values in the cingulate gyrus (β = 0.035, p_corrected_ = 0.013) and parahippocampal (β = 0.032, p_corrected_ = 0.014) part of cingulum and in the uncinate fasciculus (β = 0.029, p_corrected_ = 0.018).

### Permutation analysis

NETRIN1 was found to be individually significant in association with the following white matter tracts: superior longitudinal fasciculus as measured by FA; superior and inferior longitudinal fasciculus and anterior and superior thalamic radiations as measured by MD. In addition, NETRIN1 was solely significantly associated with thalamic radiations globally, as measured by MD. Therefore, we additionally performed a circular genomic permutation analysis in order to test whether the significant effect of NETRIN1 on these five white matter tracts and one tract category was not due to chance (table 5).

**Table 5.**
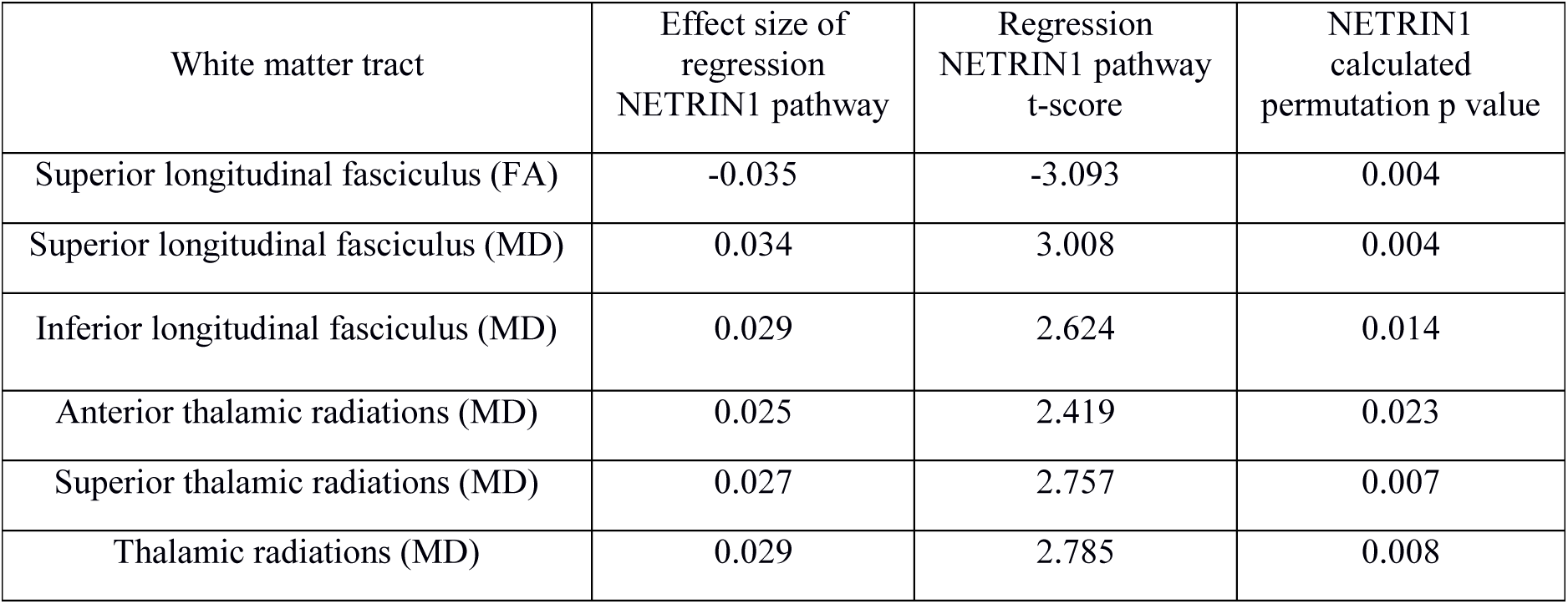
Permutation results for NETRIN1-PRS at PRS threshold 0.5 on 5 significant white matter tracts and one significant tract category.

## Discussion

In the present study, we aimed to investigate whether PRS calculated from the NETRIN1 signalling pathway are significantly associated with WM integrity after adjusting for genomic-PRS in more than 6,000 individuals. We found significant differences in white matter integrity in both NETRIN1-PRS and genomic-PRS, for both FA and MD values. Regarding FA values, for NETRIN1-PRS, a significant association was observed in the superior longitudinal fasciculus. NETRIN1-PRS were significantly associated with higher generalised thalamic radiations as measured by MD, as well as higher MD in the superior and inferior longitudinal fasciculus, and the anterior and superior thalamic radiations. Genomic-PRS were also significantly associated with FA and MD values in several tracts.

Our main finding was of an MD increase in the thalamic radiations tract category. Thalamic radiations connect the thalamus with numerous cortical areas (33), and are connected to various cognitive processes, such as attention and wakefulness (34). Thalamocortical axons play an important role during development, as their projection from the dorsal thalamus (DT) transmit sensory information to the neocortex (33). Thalamic radiations have previously been linked to MDD in numerous studies. For instance, a decrease in FA was found in the TR subset in a large UKB sample comparing 335 MDD patients with 754 healthy individuals (20). This tract subset was also found to be significantly associated with higher PRS, indicating that there is a link between the sets of tracts and a potential genetic predisposition to MDD (35).

NETRIN1, and its receptor DCC, one of the genes in the NETRIN1-pathway, have been previously implicated in thalamic axonal growth. NETRIN1 promotes growth of thalamocortical axons by binding to and activating DCC, which is expressed in the dorsal thalamus. Moreover, NETRIN1 has been shown to enhance axonal growth in explants of the DT, as well as providing guidance from the DT to the cortex (33). It has also been found that serotonin, which is highly implicated in MDD, modulates the effect of NETRIN1 on embryonic thalamocortical axons (33; 34; 36). The active involvement of NETRIN1 in thalamocortical axonal growth, therefore, may explain our findings, and further confirms that there is a potential link between a biological pathway and specific neurobiological markers in MDD.

Several other tracts also showed a significant association of FA (individually in forceps major and minor and uncinate fasciculus, and in global measures of FA and projection fibres) and MD (individually in cingulate part of the cingulum, parahippocampal part of cingulum and uncinate fasciculus, and in global measures of association and projection fibres) with genomic-PRS, most of which have also been previously associated with MDD. (20, 35). This evidence further confirms that there is an association between genetic predisposition to MDD and disruptions in white matter integrity for variants that lie outside the NETRIN1-DCC pathway.

The current study has several strengths and a few potential limitations. First of all, it is the largest combined genetic and neuroimaging study investigating the effect of PRS derived from a specific biological pathway on white matter integrity, to our knowledge. Moreover, our analysis consisted of a population-based sample of ambulant individuals recruited to UKB. Our findings might therefore be robust and generalizable to other samples within a certain age range, although studies such as UKB are not immune to biases associated with study participation, such as collider bias (37).

In addition to the large sample, the fact that NETRIN1-PRS are derived from only 43 genes, comprising approximately 0.215% of the genes in the whole genome (N = ∼ 20,000) suggests that the findings reported here are less likely to be due to chance, which was further confirmed by the permutation analysis carried out. The association between the NETRIN1 pathway and white matter integrity is therefore likely to reflect the importance of a specific pathway in the pathophysiology of MDD.

The NETRIN1 signaling pathway has previously been found to be implicated in MDD (14). In the current study, we were able to find specific neurobiological structural connectivity markers associated with this biological pathway. To our knowledge, the current study is the first one to note an association between PRS derived specifically from the NETRIN1 signaling pathway and several white matter tracts in a large genetic and neuroimaging dataset. This indicates that these brain structures may be involved in the manifestation of genetic risk of MDD and ultimately the aetiology of the disorder.

